# REM sleep predicts reductions in pathophysiological daytime basal ganglia-cortical circuit activity in Parkinson’s disease

**DOI:** 10.1101/2025.05.07.652676

**Authors:** Jin-Xiao Zhang, Clay Smyth, Hanna Cattan-Hayat, Md Fahim Anjum, Yue Leng, Andrew D. Krystal, Philip A. Starr, Simon Little

## Abstract

Sleep disturbances have been shown to be intimately and bidirectionally related to disease progression across a wide range of neurodegenerative disorders including Parkinson’s disease (PD) and Alzheimer’s disease. However, the precise neurophysiological mechanisms relating abnormal sleep to aberrant daytime network activity that accelerates disease progression has yet to be determined. We collected chronic, multi-night (*n*=40), intracranial cortico-basal recordings during sleep from a cohort of patients with PD along with paired polysomnography and morning self-reports. This revealed that longer duration (and shorter latency) of rapid eye movement (REM) sleep predicted reduced daytime resting beta (13-30 Hz) activity and cortico-basal functional and effective connectivity, features established to be pathophysiological in PD. Within REM sleep, stronger cortical delta activity specifically predicted reduced pathophysiological cortico-basal neural network features. Additionally, REM delta power significantly predicted greater self-reported morning alertness. These findings highlight a potentially protective role of REM sleep in cortico-basal network health in PD and daytime subjective experience, representing a potential target for closed loop neuromodulation therapies to impact neurodegenerative disease progression.

---

Sleep disturbance is associated with a broad range of neurodegenerative diseases, including Parkinson’s disease (PD), Alzheimer’s disease (AD), vascular dementia (VaD), frontotemporal dementia (FTD), dementia with Lewy bodies (LBD), and Huntington disease (HD) (Musiek and Holtzman 2016; Shi et al. 2018; Porter, Buxton, and Avidan 2015). Increasing evidence has shown that sleep disturbances happen early in the disease course and may precede the onset of more typical symptoms of neurodegenerative diseases (Schenck, Boeve, and Mahowald 2013; He, Hu, and Jiang 2020; Irwin and Vitiello 2019; Rothman and Mattson 2012). Importantly, recent evidence suggests that sleep disturbances could also contribute to the pathogenesis of neurodegenerative diseases and accelerate disease progression in a bidirectional relationship (Shokri-Kojori et al. 2018; J. E. Owen and Veasey 2020; J. Owen et al. 2021; Ju, Lucey, and Holtzman 2014). The protective role of sleep in neurodegenerative diseases encompasses both cellular/proteomic mechanisms as well network synaptic homeostasis, in which sleep regulates synaptic plasticity and connectivity (Tononi and Cirelli 2014; Kharas et al. 2024).

Prior sleep research, with a particular focus on non-rapid eye movement (NREM), has uncovered many cellular and molecular mechanisms linking sleep disturbance to neurodegenerative diseases, inc. putative disruption of glymphatic clearance (Musiek and Holtzman 2016; C.-W. Chen et al. 2023). However, the neurophysiological mechanisms by which sleep impacts brain network health and connectivity in neurodegenerative conditions, including PD, remain less understood. Given the high prevalence of sleep disturbances in neurodegenerative diseases, elucidating a neurophysiological link between sleep and network health will identify potential interventional targets during sleep for preventing and slowing neurodegenerative diseases. Human sleep is broadly classified into alternating stages of NREM sleep and rapid eye movement (REM) sleep. Electroencephalogram (EEG) in NREM sleep is hallmarked by larger-amplitude lower-frequency activity, especially during the deep NREM stage (i.e., N3) (Berry et al. 2017). During REM sleep, EEG transitions to a low-amplitude mixed-frequency desynchronized activity. Although the neurophysiological activity during REM has generally higher frequencies compared to NREM, recent evidence has shown the existence of slow-wave (i.e., delta, 1-4 Hz) activity during REM sleep in both humans and mice, particularly in the frontal-central region (Bernardi et al. 2019; Funk et al. 2016). The role of these REM slow waves has yet to be established.

Animal studies have demonstrated the benefits of NREM slow-wave sleep for neural network health, associated with glymphatic clearance (Xie et al. 2013), synaptic renormalization (Tononi and Cirelli 2014), and neuroinflammation reduction (Wisor, Schmidt, and Clegern 2011; Zielinski and Gibbons 2022). In studies of human neurodegenerative diseases, NREM slow-wave activity during sleep is associated with slower motor progression in PD (Schreiner et al. 2019; J. Chen et al. 2024), improved memory function, and slower accumulation of pathological beta-amyloid in AD (Wunderlin et al. 2020; Lee et al. 2020; Ju et al. 2017). Previous studies have focused on NREM sleep when investigating the role of slow-wave activity in neurodegenerative diseases. However, recent discovery of slow-wave activity during REM sleep opens the question of whether REM delta activity may also impact brain health in neurodegenerative diseases (Bernardi et al. 2019; Funk et al. 2016), including PD (Mizrahi-Kliger et al. 2018; Zahed et al. 2021).

PD is a common and increasingly prevalent neurodegeneration condition intimately related to sleep disturbance (Zahed et al. 2021; Lima 2013). Over 80% of REM sleep behavior disorder (RBD) patients develop PD or neurodegenerative dementia over a median of 14-year period (Schenck, Boeve, and Mahowald 2013; Boeve 2013), suggesting a potential role of REM sleep in the pathogenesis of PD. Moreover, up to 90% of patients with PD experience disturbed sleep, which is a major contributor to poor quality of life in this patient population (Videnovic and Golombek 2013). Relative to healthy controls, patients with PD exhibit a reduced REM percentage and prolonged REM latency, while changes in NREM percentage and latency are more modest (Zhang et al. 2020; Yong et al. 2011). A new generation of advanced sensing-enabled deep brain stimulation (DBS) devices provide a unique window to directly study the sleep neurophysiology of patients in a chronic and ecologically valid environments. Additionally, these techniques allow, for the first time, the collection of longitudinal multi-night intracranial data, that affords the opportunity to examine the relationship between overnight sleep features and next day brain network health (Gilron et al. 2021; Oehrn et al. 2024).

During daytime, PD is characterized by pathophysiological features in the cortico-basal network, most prominently excessive activity and heightened connectivity in the beta (13-30 Hz) band (Hammond, Bergman, and Brown 2007; Simon Little and Brown 2014; Uhlhaas and Singer 2006; Moran et al. 2011; Simon Little et al. 2013). Cortico-basal beta activity predicts motor symptom severity in patients with PD, indexes disease progression, and potentially indicates pathological inter-region connectivity (S. Little et al. 2012; C. C. Chen et al. 2010; Morelli and Summers 2023; Neumann et al. 2017; Stoffers et al. 2008). Given the role of sleep in subserving neural plasticity across multiple brain networks (Verweij et al. 2014; Tashjian et al. 2018; Di et al. 2024; Kaufmann et al. 2016; Huber et al. 2013), we hypothesized that sleep physiology would impact synaptic renormalization processes (Tononi and Cirelli 2014), thereby affecting connectivity across the cortico-basal network in patients with PD. Specifically, we predicted that longer sleep duration and shorter sleep latency would be associated with reduced pathophysiological resting-state beta activity and connectivity features in the basal-cortical network following sleep. Given the demonstrated neuroprotective effects of delta activity in neurodegeneration (Schreiner et al. 2019; Wunderlin et al. 2020), we anticipated that stronger delta activity during sleep would be associated with reduced pathophysiological features. Complementarily, considering the pathological role of beta activity during sleep in PD (Anjum et al. 2024), we predicted that increased beta activity during sleep would be harmful and self-reinforcing in network connections, leading to increased pathophysiological neural signals during the daytime - a “snowball” effect. Considering the established role of dysfunctional REM sleep in PD (Schenck, Boeve, and Mahowald 2013), as well as previous between-subject studies demonstrating a link between NREM sleep and disease progression (Schreiner et al. 2019; J. Chen et al. 2024), we tested the effects of both REM and NREM sleep and directly compared them in this study. We investigated the effects of REM and NREM sleep on daytime neurophysiology in patients with PD, with particular emphasis on the aspects of sleep neurophysiology which may protect against pathophysiological activity and connectivity in PD.

We conducted a within-subject, at-home, multi-night intracranial recording investigation in patients with PD, investigating naturalistic fluctuations in sleep and the impact on daytime cortico-basal networks. Participants were implanted with bilateral chronic electrocorticography (ECoG) spanning the central sulcus and sensing- and stimulation-capable electrodes in the basal ganglia (BG). This architecture supports high volumes of temporally resolute neural data, collected across multiple nights. Through this approach, we discover neural determinants of healthy REM sleep that reduce pathophysiological daytime network activities and connectivity known to be associated with worse symptoms in Parkinson’s disease.

## Results

We obtained chronic multi-night (total *n*=40) intracranial BG-cortical neurophysiological recordings (total recording hours: left device = 218.7, right device = 191.6) from four patients with PD during overnight sleep and next-morning wakefulness. Participants received bilateral surgical implantation of sensing and stimulation capable, cylindrical quadripolar electrode leads (Medtronic model 3389) into the basal ganglia, either the subthalamic nucleus (STN) or the globus pallidus internus (GPi) (Figure 1A) as part of a parent study (Oehrn et al. 2024). Additionally, four-electrode ECoG arrays were bilaterally implanted in the subdural space over the sensorimotor cortex, spanning the central sulcus (Medtronic model 0913025). Ipsilateral basal ganglia leads and ECoG arrays were connected to an investigational sensing-enabled pulse generator (Medtronic Summit RC+S model B35300R), which was secured above the pectoralis muscle and capable of chronically streaming high-resolution time domain data from both cortex and basal ganglia (Figure 1B).

**Figure 1.**
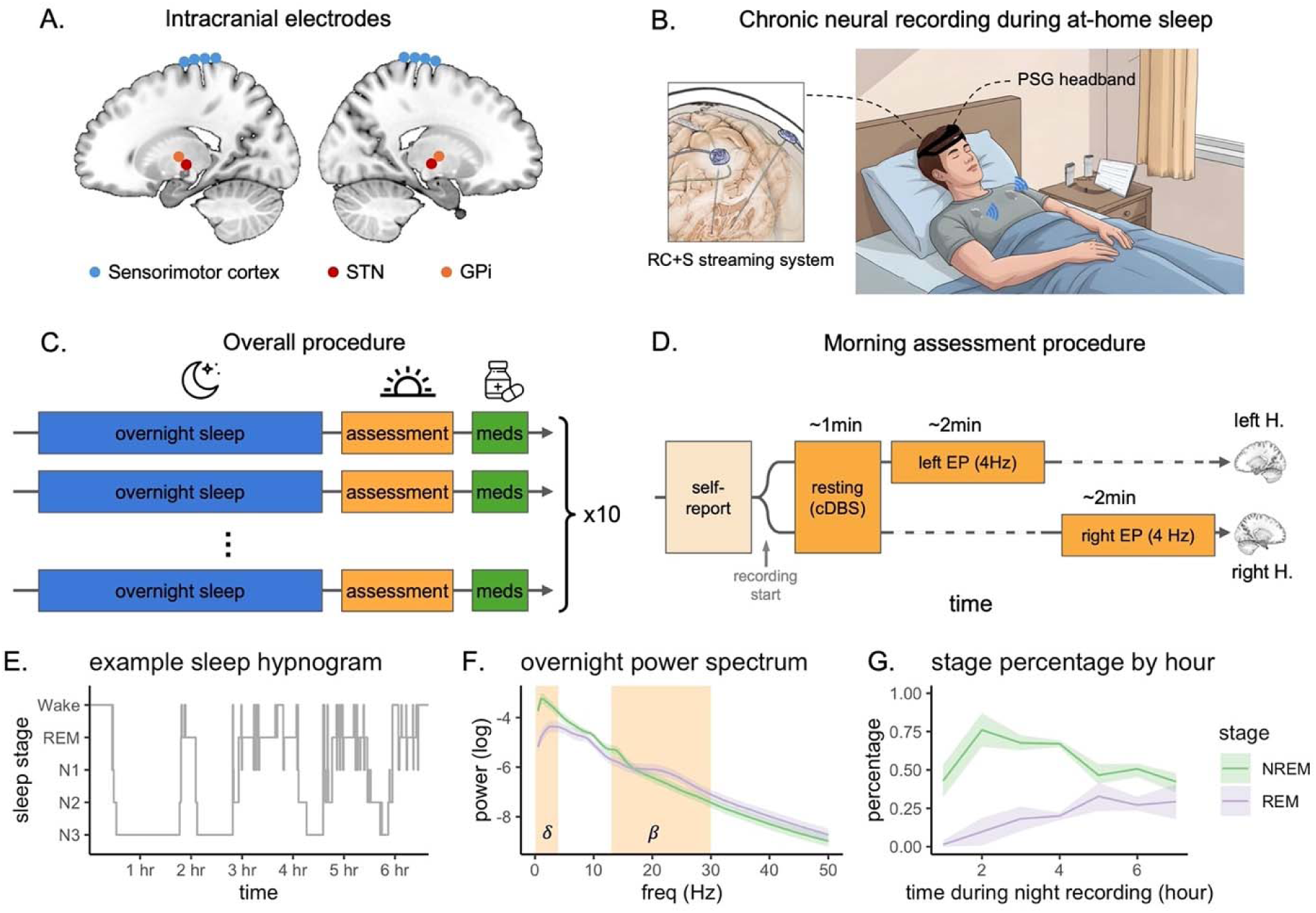
Study overview. (A) Illustrations of the placement of the intracranial RC+S electrodes in the sensorimotor cortex and subcortical basal ganglia. (B) Schematic of neural recording setup during sleep. Data were concurrently recorded using the intracranial RC+S streaming system and a portable polysomnography (PSG) system (the DREEM2 headband). (C) An overview of the multi-night-morning protocol containing an overnight recording session and a morning recording session for approximately 10 days. Morning sessions took place after wake-up and before dopamine medication use. (D) The procedure of the morning assessment sessions. The morning sessions contained self-report surveys, a resting-state recording session with constant DBS (cDBS), and an evoked potential (EP) session in each hemisphere. Dotted lines indicate absence of DBS in the corresponding hemisphere. (E) The sleep hypnogram of a sample night recording. Each 30s epoch during overnight recordings was scored into wake, REM, N1, N2, or N3 stage. (F) Power spectra of the intracranial cortical data over cortical channels in REM and NREM sleep in all participants. (G) The percentage of REM and NREM during each hour segment of the overnight recordings in all participants.

Nighttime intracranial recordings were complemented by portable extracranial polysomnography (PSG, DREEM2) (Figure 1B). Sleep stages were scored into REM or NREM 1-3 stages by a validated automated algorithm (Arnal et al. 2020; González et al. 2024; Ravindran et al. 2025). Each morning, after awakening and before taking medications, participants completed standardized subjective self-report and objective neurophysiological evaluations (Figure 1C). The neurophysiological assessment included resting-state recordings to assess spectral features and functional connectivity in the BG-cortical circuit, as well as effective connectivity in a BG → cortex evoked potential (EP) paradigm (Figure 1D). We used linear mixed-effects (LME) models to perform within-patient evaluations on how nighttime sleep attributes predicted pathophysiological daytime BG-cortical activity and connectivity. To directly compare features of REM and NREM sleep and to reduce the number of tests, we included them as predictors in the same LME models. These LME models tested separate hypotheses about sleep features (duration vs. power) and neural network health outcomes (resting vs. evoked state), and thus did not undergo further multiple-test corrections. Following our hypotheses, in the ‘sleep metric’ LME models, we tested the effects of the duration and latency of REM and NREM sleep on morning BG-cortical pathophysiological activity and connectivity. In ‘spectral power’ LME models, we tested the effects of delta and beta power during REM and NREM sleep on the same morning features. In exploratory analyses, we also tested all power bands from 1-90 Hz in separate LME models (Supplementary Materials). All LME models converged well.

### Overnight sleep and sleep neurophysiology

Each participant’s intracranial cortical and subcortical signals were recorded for ∼10 (generally consecutive) nights. Participants received their standard therapeutic (continuous) DBS throughout the study period (nighttime and daytime, except the EP paradigm) and this was held constant for the duration of the recordings. On average, participants slept for 6.1±1.1 hours each night and achieved 79.9% (SD=7.8%) sleep efficiency. According to hypnograms (Figure 1E), participants spent 22.0% (SD=8.3%) of their sleep during the REM stage and 68.8% (SD=8.5%) during NREM (N2 and N3 combined for later analysis) stages. As expected, the REM sleep percentage was greater (*t*(302) = 3.50, *p* < .001) and the NREM sleep percentage (*t*(408) = -6.89, *p* < 10^-5^) was smaller in the latter half of the night than the first half (Figure 1G). We calculated the spectral power of cortical ECoG channels during REM and NREM stages (Figure 1F; for all stages, see Supplementary Figure 1A). Similar to healthy individuals, our PD participants typically had stronger power in the delta band and smaller power in higher frequency bands (beta and above) during NREM than REM sleep.

### REM sleep predicts resting-state pathophysiological features in BG-cortical network

During the morning resting-state recording, intracranial signals from BG and sensorimotor cortex were streamed in both hemispheres while participants received their therapeutic constant DBS (Figure 1D). A beta-band power peak was observed in the cortical channels but was suppressed in the BG under continuous DBS (Supplementary Figure 1B) (Binns et al. 2025), which directed our following analysis to focus on the cortical channels. As expected, bursts of beta oscillation were observed during the resting recording period. The density and magnitude of beta bursts differed following nights of short REM sleep vs. long REM sleep (Figure 2A). Within-individual median splits of REM duration demonstrated that nights with longer REM duration were associated with less resting cortical beta power the next morning (Figure 2B). Using LME models, we tested whether the ‘sleep metric’ attributes (duration, latency) or the ‘spectral power’ attributes (delta power, beta power) of REM or NREM sleep predicted the morning resting-state beta power. In the ‘sleep metric’ model, REM sleep duration was a negative predictor of resting beta power, *t* = -2.77, *p* = .007 (Figure 2C). Furthermore, in the ‘spectral power’ model (Figure 2D), the delta power during REM sleep was also a negative predictor of resting cortical beta power, *t* = -2.27, *p* = .027. In addition, beta power during NREM sleep positively predicted resting beta power, *t* = 2.04, *p* = .047. Although the NREM sleep duration and delta power showed the same correlative trend as their REM counterparts in the LME models, they were not significant predictors (for results of testing NREM and REM predictors separately and additional frequency bands, see Supplementary Figure 2). These results suggest that longer REM sleep and stronger delta activity during REM are associated with less pathological cortical beta oscillations in the following morning.

**Figure 2.**
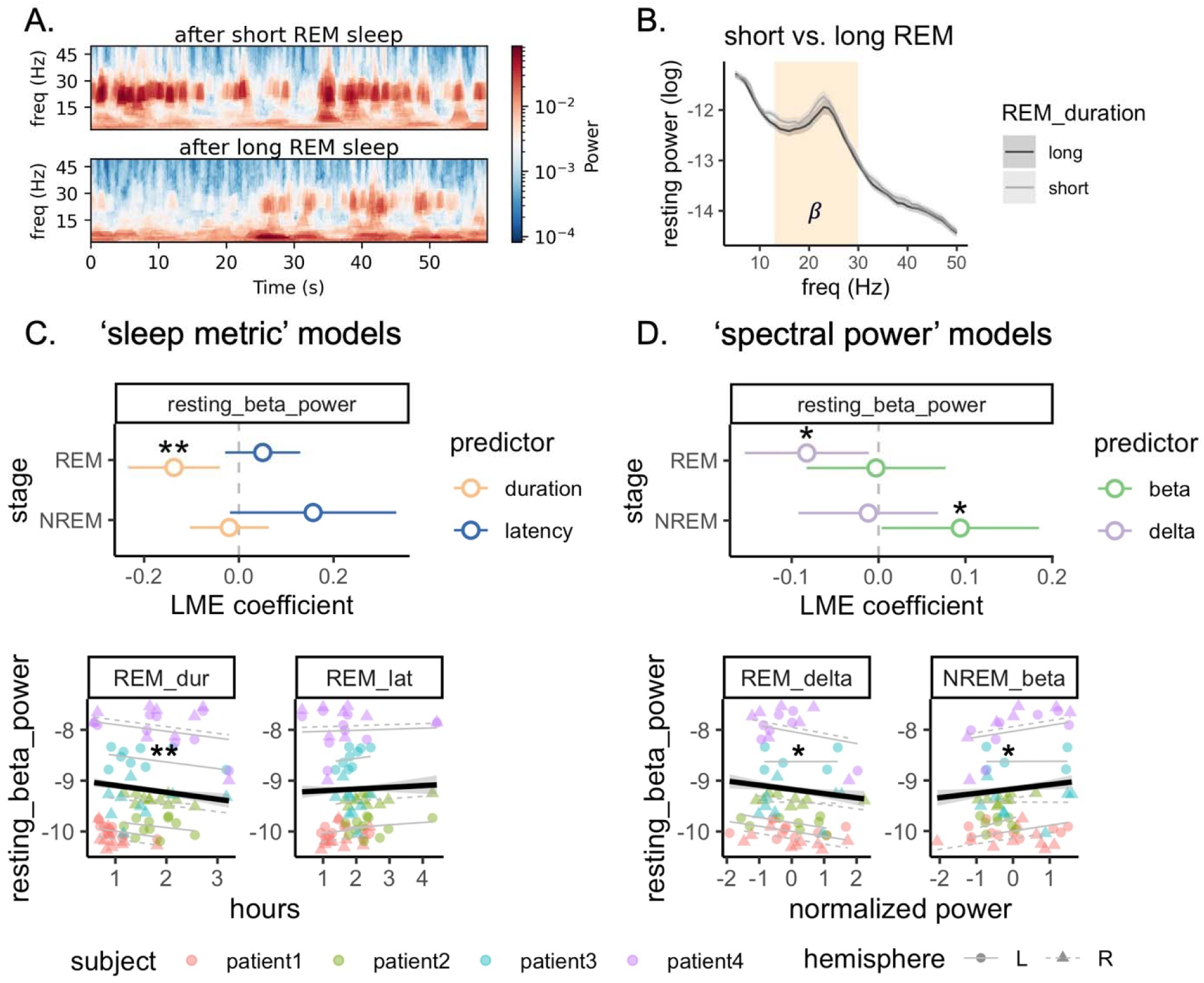
REM sleep predicts next-morning resting-state cortical beta power. (A) Morning resting-state power spectrogram following an example night of short REM sleep (0.7 h) vs. an example night of long REM sleep (3.2 h) in one participant. (B) The group average morning resting-state power spectrum following nights of long REM duration vs. nights of short REM duration (with median splits in each participant). The highlighted area shows the frequency range of beta bands (13-30 Hz). (C) The results of linear mixed-effect (LME) models of sleep metric predictors (duration and latency) in prediction of resting-state beta power. The top panel shows the LME coefficients of duration and latency of REM and NREM sleep with 95% confidence intervals. The bottom panel shows the scatter plot of resting-state beta power against REM duration and latency. (D) The results of LME models of sleep spectral power predictors (cortical beta and delta) in prediction of resting-state beta power. The top panel shows the LME coefficients of delta and beta power during REM and NREM sleep with 95% confidence intervals. The bottom panel shows the scatter plot of resting-state beta power against REM beta and delta. In scatterplots, model predictions were shown at both the hemisphere level and the group level. REM_dur: REM duration; REM_lat: REM latency. Asterisks indicate statistical significance in LME models: * = *p*<.05, ** = *p*< .01.

We also calculated the BG-cortical coherence in beta frequency range to index the functional connectivity between motor cortex and BG. LME models showed that REM duration was a negative predictor of the BG-cortical coherence at beta frequencies, *t* = -2.58, *p* = .014, and so was REM latency, *t* = -3.88, *p* < .001, suggesting that longer and later REM sleep was associated with less pathological BG-cortical hyper-connectivity the next morning.

### REM sleep predicts BG-to-cortical effective connectivity

During the later section of the morning session (Figure 1D), we used EPs to causally assess effective connectivity between BG and motor cortex. Specifically, stimulation pulses of 4mA were delivered to BG via the implanted DBS leads at a rate of 4 Hz (i.e. 250 ms per epoch) while cortical field potentials were sampled at 1000 Hz. This procedure was implemented for 2 minutes in each hemisphere at a time, with the contralateral DBS off.

The evoked responses in the motor cortex consisted of an initial early response (<10 ms) which likely reflects antidromic activation of the hyperdirect pathway (Ashby et al. 2001; Baker et al. 2002; W. Chen et al. 2020), and a later decaying oscillatory response (>10 ms) at primarily a beta frequency around 20 Hz which likely reflects orthodromic activation of BG projections to the cortex, plus beta resonance (Baker et al. 2002; MacKinnon et al. 2005; Eusebio et al. 2009) (Figure 3A, 3B). Increased evoked response in the beta frequency range supports that circuit connectivity is naturally resonant for pathological oscillatory activity in the beta band. We assessed attributes of the EP signal to index pathophysiological hyperconnectivity in the BG-motor cortex circuit (Campbell et al. 2022). Specifically, we calculated the peak-to-peak amplitude in the initial early window (0 to 10 ms), the peak-to-peak amplitude in the later main window (10 to 100 ms), plus the beta power in the whole orthodromic window (10 to 200 ms) of the evoked response (longer window to enable a FFT with sufficient frequency resolution). In the ‘sleep metric models (Figure 3C), we found that REM duration negatively, and REM latency positively, predicted the main-window BG-cortex evoked amplitude (duration: *t* = -2.57, *p* = .012, latency: *t* = 2.57, *p* = .012) and the whole-window BG-cortex induced beta power (duration: *t* = -2.47, *p* = .016, latency: *t* = 2.16, *p* = .034). Neither duration feature was a predictor of the early-window evoked amplitude (duration: *t* = -0.60, *p* = .550, latency: *t* = 0.49, *p* = .628). In the ‘spectral power’ models (Figure 3D), we found that REM delta power negatively predicted the main-window BG-cortex evoked amplitude (*t* = -2.16, *p* = .036) and whole-window BG-cortex induced beta power (*t* = -3.01, *p* = .004). Again, these power features were not predictors of the early-window evoked amplitude (*t* = -0.86, *p* = .391). Sleep beta power, in REM or NREM, generally showed a positive trend in predicting EP metrics, but did not reach statistical significance (*p*’s > .20). In models which tested REM and NREM together, NREM sleep attributes were not significant predictors, *p*’s > .05 (for results of testing NREM and REM predictors separately and additional frequency bands, see Supplementary Figure 3-5). Altogether, these results suggest that longer duration of, quicker entry into, and stronger delta activity in REM sleep predicted less pathological basal-cortical effective connectivity the following morning.

**Figure 3.**
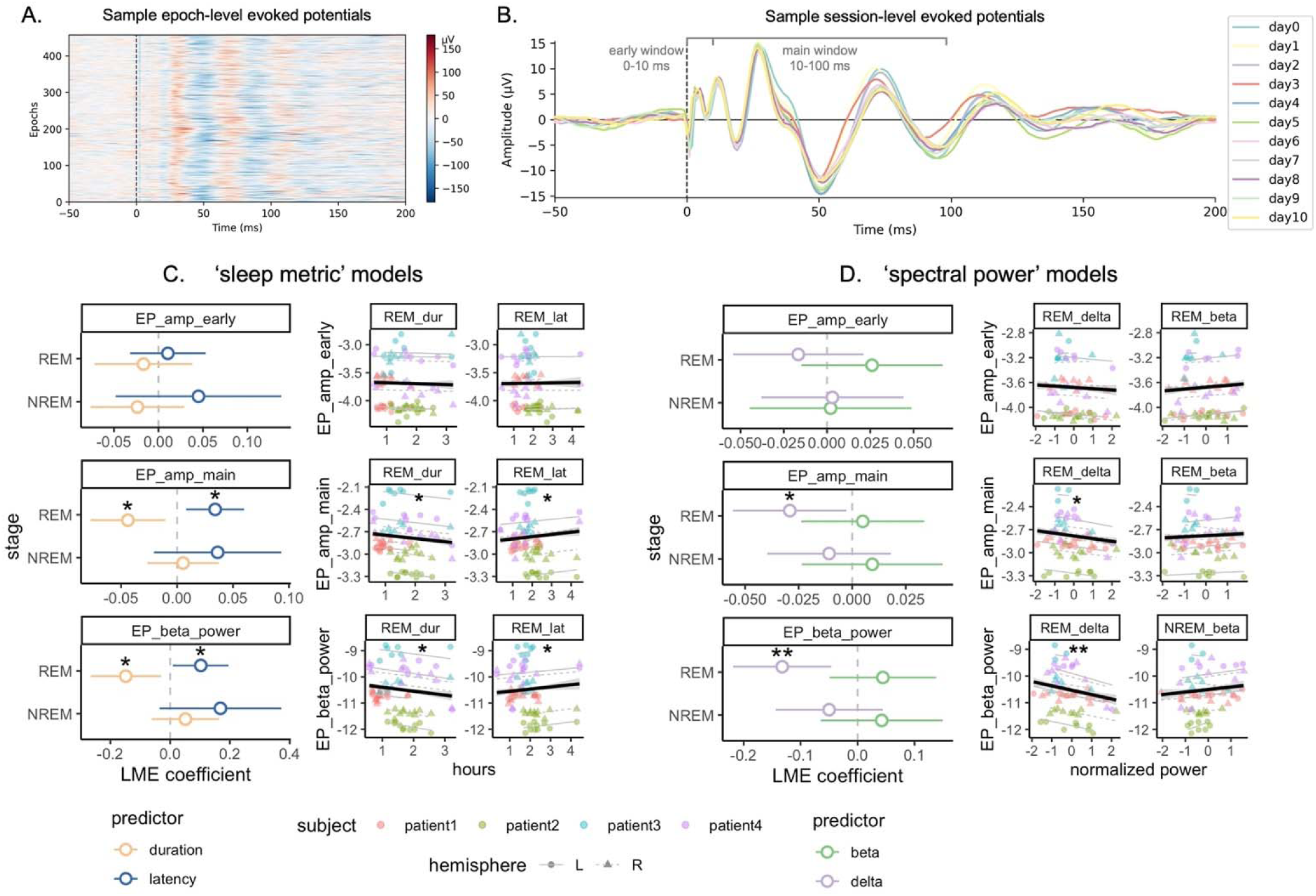
REM sleep predicts next-morning evoked potentials (EP) in the basal ganglia-cortical circuit. (A) Trial-level cortical EP data of an example morning session. Each epoch spans from -50ms to 200 ms. (B) Session-level averaged waveform of cortical EP of an example participant. At the trial level, we calculated the peak-to-trough amplitudes in an early window (0–10 ms) and a main window (10–100 ms), and the beta power in the entire window (10–200 ms) and then averaged across epochs. (C) The results of linear mixed-effect (LME) models of sleep metric predictors (duration and latency) in prediction of EP metrics. (D) The results of LME models of sleep spectral power predictors (cortical beta and delta) in prediction of BG-cortex EP metrics. In (C) and (D), the left panels show the LME coefficients with 95% confidence intervals and the right panels show the scatterplots of outcome variables against REM predictors with model predictions. ep_amp_ealry: peak-to-trough amplitude of EP in the early window; ep_amp_main: peak-to-trough amplitude of EP in the main window; ep_beta_power: beta power of the EP in the whole window; REM_dur: REM duration; REM_lat: REM latency. Asterisks indicate statistical significance in LME models: * = *p*<.05, ** = *p*< .01.

### Detection of delta waves and beta bursts during sleep

Given the apparent importance of sleep delta activity shown above, we further evaluated individual delta waves during REM and NREM sleep. As the metric of delta power naturally combines both the amplitude and frequency of the occurrence of delta waves, we further separately analyzed the amplitude and occurrence rate of individual waves, and compared these results with overall delta power, in terms of predictive magnitude. Following prior research (Bernardi et al. 2019), we detected delta half-waves in the cortex during sleep using a consecutive zero-crossings algorithm. The detected delta half-waves were present in both REM and NREM sleep stages, with greater rate of occurrence in NREM (N3 and N2) than REM, *F*(2, 425) = 62.7, *p* < 10^-5^ (Figure 4B). The waveform morphologies showed a non-sinusoidal shape, with the average waveform containing a sharp peak (Figure 4A). As expected, the average peak amplitude of delta waves was larger in NREM (N3 and N2) than REM sleep, *F*(2, 425) = 132.8, *p* < 10^-5^ (Figure 4C). We further tested, in separate LME models, whether the average peak amplitude or occurrence rate of these delta waves predict next-morning pathophysiological features (resting beta power, EP amplitude, and EP beta power). As the outcome variables and the predictors are in different scales, we compared the standardized coefficients (*t*-stat) in the LME models. Compared to the coefficients of overall delta power (mean *t* = -1.51), average peak amplitude of delta waves had greater or comparable coefficients (mean *t* = -2.51) while their occurrence rate generally had smaller coefficients (mean *t* = -0.86) (Figure 4D), underscoring the importance of the peak amplitude of these delta waves on morning pathophysiological features. A *t*-test showed that the attributes of delta waves in REM had greater-sized coefficients than those of NREM stages (N3 and N2), *t*(25) = 3.23, *p* = .003, highlighting the role of REM-specific delta waves.

**Figure 4.**
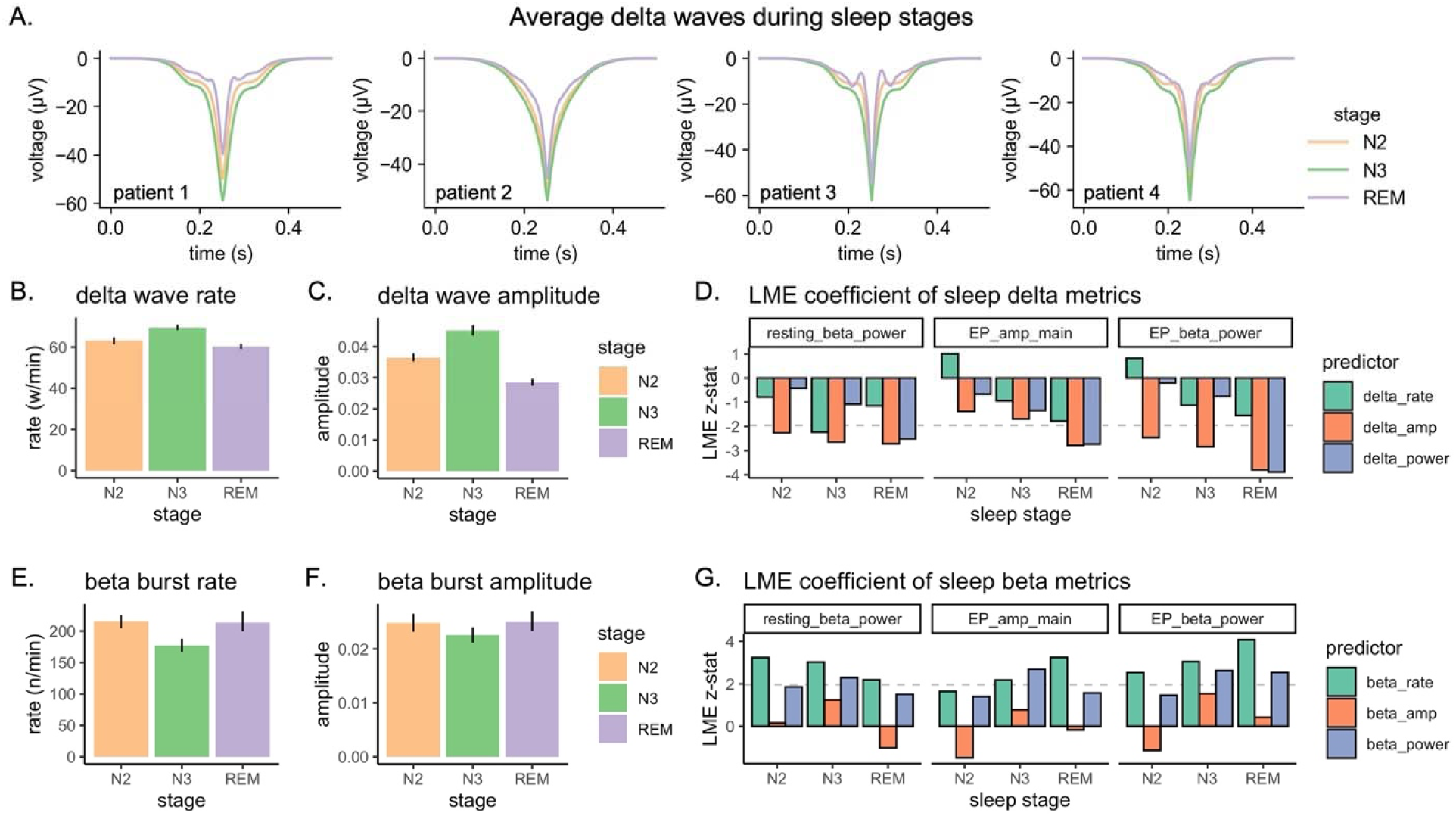
Detection of delta waves during sleep. (A) Averaged negative delta half-waves detected during N2, N3, and REM stages in each patient. The individual delta half-waves were aligned at their largest negative peak and averaged accordingly. Occurrence rate (B) and amplitude (C) of detected delta waves across sleep stages. (D) Standardized LME coefficients of delta wave metrics across sleep stages in prediction of morning pathophysiological features. Dashed lines indicate LME *t-stat* = -2. Occurrence rate (E) and amplitude (F) of beta bursts across sleep stages. (G) Standardized LME coefficients of beta metrics across sleep stages in prediction of morning pathophysiological features. Dashed lines indicate LME *t-stat* = 2.

The aforementioned results (Figure 2D and 3D) showed that sleep beta power was associated with increased morning pathophysiological neural signals. For comparison, we also detected beta bursts during sleep and calculated their amplitude and occurrence rate. We observed that beta bursts had a higher occurrence rate (*t*(137) = 3.91, *p* < .001) and a greater amplitude (*t*(137) = 2.00, *p* = .047) in REM than in N3 (Figure 4E and 4F). When tested in separate LME models to predict morning pathophysiological features (Figure 4G), in contrast to our delta findings, beta burst occurrence rate had greater standardized coefficients (mean *t* = 2.79) than overall beta power (mean *t* = 1.99) while the beta burst amplitude showed smaller coefficient (mean *t* = 0.03), highlighting the uniquely pathological role of beta bursting rate during sleep. The coefficients of these beta bursts attributes were not significantly different between REM and NREM stages, *t*(25) = 0.03, *p* = .973.

### Timing effect of REM sleep

Our analyses above showed a critical role of REM sleep in down-regulating pathophysiological BG-cortical activity and connectivity in PD. One question is whether this protective role is influenced by the temporal structure of NREM and REM sleep. To answer this question, we calculated the relative proportion of time spent in NREM versus REM sleep (Figure 1G) and the delta power in NREM versus REM throughout the sleep course (Figure 5A). Along with the percentage of REM sleep, the delta power during REM sleep also increased moderately over the sleep course, *t* = 2.31, *p* = .025 (Figure 5B). In order to test the role of REM and NREM sleep individually, we tested their effects in prediction of next-morning pathophysiological features in separate LME models for each hour segment.

**Figure 5.**
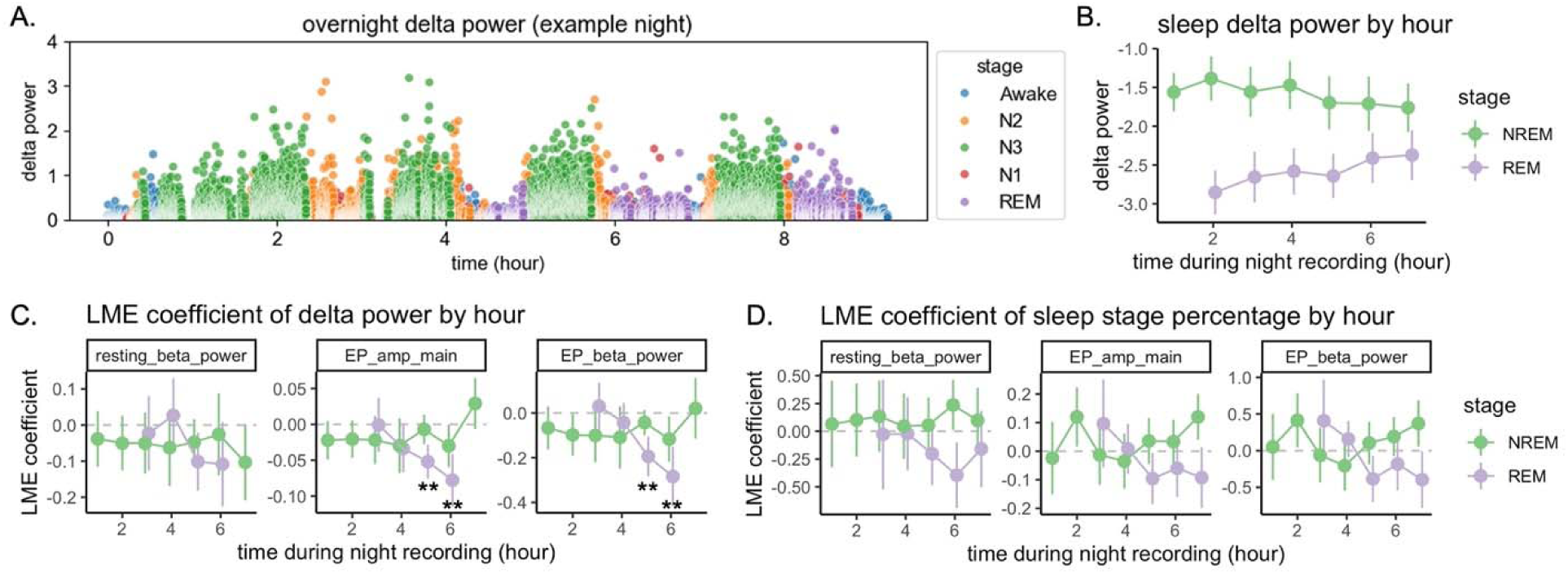
Timing effect of REM sleep over the sleep course. (A) Delta power over the sleep course of an example night. (B) Average delta power in REM and NREM sleep during each hour segment over the sleep course across participants. (C) LME coefficients of REM and NREM delta power by each hour segment in prediction of morning pathophysiological features. Predicting EP amplitude in the main window: REM delta was a negative predictor at hour 5 (*t* = -4.28, FDR corrected *p* = .002) and hour 6 (*t* = -4.18, FDR corrected *p* = .002), and NREM delta was not a significant predictor at any hour. Predicting EP beta power: REM delta was a negative predictor at hour 5 (*t* = -4.29, FDR corrected *p* = .002) and hour 6 (*t* = -4.18, FDR corrected *p* = .002), and NREM delta was not a significant predictor at any hour. All other LME tests were non-significant after FDR correction, *p*’s >.05. (D) LME coefficients of REM and NREM duration percentage by each hour segment in prediction of morning pathophysiological features. All LME tests were non-significant after FDR correction, *p*’s >.05. Error bars indicated 95% confidence intervals. In (C) and (D), asterisks indicate statistical significance after FDR correction: * = *p*<.05, ** = *p*< .01.

Delta power during early REM sleep was a null predictor of the effective connectivity indices (i.e. main-window amplitude and beta power of EP). However, the predictive effect of REM delta grew stronger and became a significant negative predictor for later hours, FDR corrected *p*’s <.01 (Figure 5C). NREM delta power, in contrast, remained a null or a less strong negative predictor, and did not show a clear temporal trajectory throughout the course of sleep. This negative temporal effect for NREM demonstrates that the comparatively stronger effect of delta during REM than NREM is not simply due to the closer temporal proximity of REM sleep to the morning neurophysiological assessments. Furthermore, a similar temporal trend was observed when using the REM percentage per hour to predict morning pathophysiological features, although the statistical tests did not survive multiple test corrections (Figure 5D). These results suggest a timing effect of the protective role of REM sleep (especially its delta activity) in maintaining a healthy profile of BG-cortical circuit. The effect of REM sleep was more pronounced during the later hours of sleep.

### REM delta activity predicts subjective morning alertness

Following awakening, participants reported their subjective sleepiness using Karolinska Sleepiness Scale (Figure 6A) and “restedness” from sleep using the Consensus Sleep Diary (Figure 1D, 6B) (Kaida et al. 2006; Carney et al. 2012). These two self-report measures were negatively correlated, Pearson’s *r* = -0.56, *p* = .003. They reflected different elements of morning alertness and exhibited dissimilar distributional shapes (Figure 6A bottom left, Figure 6B bottom left). Consistent with our neurophysiological findings of delta activity being beneficial and beta activity being pathological to waking network health, we observed that stronger REM delta power (*t* = -2.21, *p* = .031) and smaller NREM beta power (*t* = 2.10, *p* = .040) were associated with lower self-reported sleepiness scores in the morning (Figure 6A bottom right). Further, stronger REM delta was also associated with greater sense of restedness after sleep, *t* = 2.56, *p* = .013 (Figure 6B bottom right). Moreover, the self-report restedness was inversely associated with the BG-cortical effective connectivity outcomes (early-window EP amplitude: *t* = -2.84, *p* = .007; main-window EP amplitude: *t* = -2.60, *p* = .013; whole-window EP beta power: *t* = -2.76, *p* = .008; LME models).

**Figure 6.**
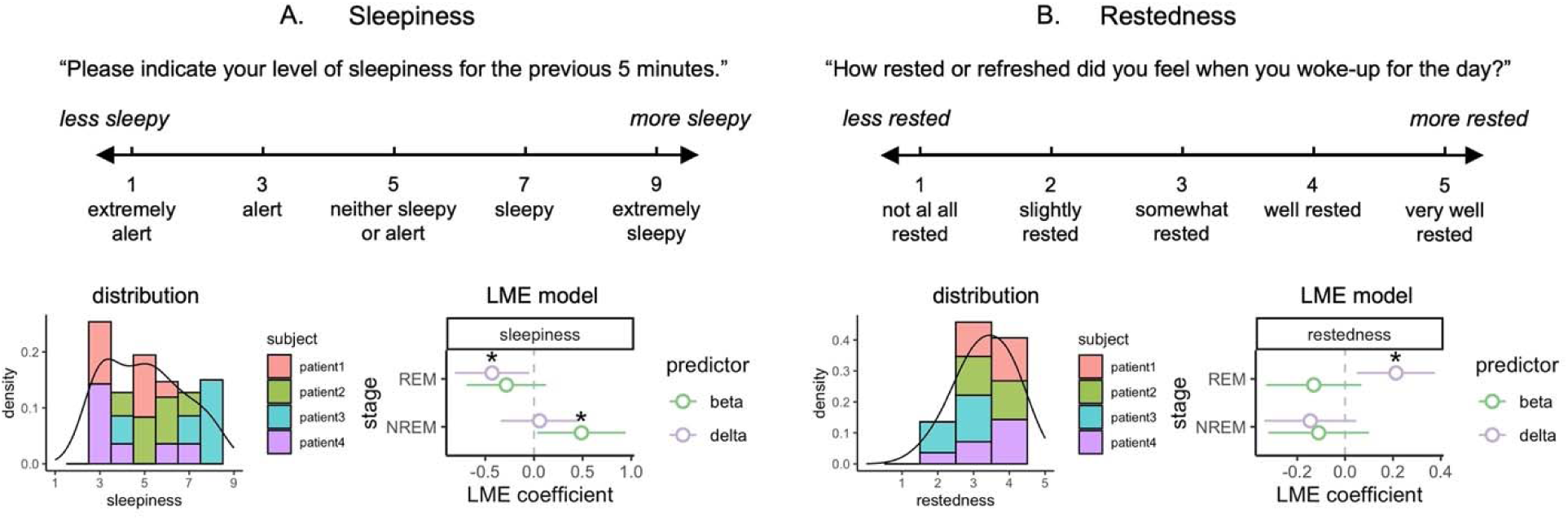
Sleep neural activity predicts morning self-report alertness. (A) Sleepiness measured by Karolinska Sleepiness Scale. Top: questionnaire instruction and the 9-point Likert scale. Bottom left: distribution of self-report sleepiness. Bottom right: results of linear mixed-effect (LME) models in prediction of sleepiness. (B) Restedness measured in Consensus Sleep Diary. Top: questionnaire instruction and the 5-point Likert scale. Bottom left: distribution of self-report restedness, larger number means more rested or refreshed upon awakening. Bottom right: results of LME models in prediction of restedness. Asterisks indicate statistical significance in LME models: * = *p*<.05, ** = *p*< .01.

## Discussion

In this study, we followed PD patients for multiple nights of sleep and subsequent mornings while recording intracranially from the sensorimotor cortex and the basal ganglia. We assessed the pathophysiological neural network features related to motor symptoms in PD (i.e., resting beta activity, plus functional and effective BG-cortical connectivity) as well as self-reported subjective states in the morning following sleep. We showed that longer REM duration and shorter REM latency predicted reduced pathophysiological features the next morning. When evaluating within REM sleep, we found that cortical delta power predicted a reduction in these same morning pathophysiological features. We further detected individual delta waves during REM and NREM sleep to demonstrate that the amplitude of these delta waves was the major mechanism to predict reduced pathophysiological features. Our timing analysis showed that the protective effect of REM sleep grew stronger across the sleep course. In contrast, the effect of NREM sleep was relatively steady and less pronounced than REM sleep throughout the night. Finally, REM delta activity was also found to be associated with reduced self-reported morning sleepiness, a highly negatively impactful symptom in PD (Knie et al. 2011).

### REM vs. NREM sleep in relation to neurodegeneration

It is notable that REM sleep is here more strongly related to next-morning pathophysiological neural network signatures of PD than NREM sleep. In our LME models, several REM parameters (e.g., longer duration, shorter onset latency, and stronger delta activity) significantly predicted reduced next-morning pathophysiological neural network signatures of PD. The NREM counterparts were not significant predictors. Although abnormalities in REM sleep (i.e., REM sleep behavior disorder) are a strong precursor of PD (Schenck, Boeve, and Mahowald 2013), most prior studies in the field have focused on NREM sleep when researching PD pathophysiology / progression (Schreiner et al. 2019; J. Chen et al. 2024) and other neurodegenerative diseases (Wunderlin et al. 2020; Lee et al. 2020; Ju et al. 2017). Our results are consistent with the emerging findings implicating delayed REM onset in disease progression of AD(Jin et al. 2025; Sharon et al. 2025).

Prior research has shown that NREM sleep is linked to glymphatic clearance in neurodegenerative diseases (Xie et al. 2013). Our results suggest that REM sleep may compensate for network health with a unique mechanism independent of the glymphatic clearance, likely related to modulation of plasticity and synaptic homeostasis (Tononi and Cirelli 2014). This suggests two distinct mechanisms linking sleep to neurodegeneration in PD and possibility other conditions including AD: 1) NREM related delta waves linked to glymphatic clearance of protein aggregates, and 2) REM-related network remodeling through plasticity mechanisms to renormalize synaptic homeostatic balance. Some prior evidence indicates that REM sleep downscales synaptic connectivity (e.g. Hebbian synaptic depression) more markedly than NREM sleep (Niethard, Burgalossi, and Born 2017; Grosmark et al. 2012; Placidi et al. 2013). Therefore, REM sleep may be particularly well-suited to combat circuit hyperconnectivity. Mechanistically, synapses in specific circuits and cell types may be primed for plastic changes during REM sleep, coincident with delta waves in REM sleep inducing hyperpolarization of ensembles of cortical neurons (Placidi et al. 2013; Renouard et al. 2022; Muheim and Frank 2025). These events may alter statistical correlations of cortical neural action potentials with afferent activity. Together, in health these events may drive Hebbian long-term-depression in over-active synapses (Feldman 2012) and provide the underlying mechanism for homeostatic correction against pathological hyperconnectivity observed in PD, and potentially other neurodegenerative disorders. This pathological “snowballing” effect, whereby abnormal sleep reinforces aberrant network plasticity changes and hyperconnectivity during the daytime, represents a significant target for responsive neurostimulation techniques.

It is also worth noting that when tested separately, the LME coefficients of NREM and REM delta activities trended in the same direction (Figure 4D, Supplementary Figure 2, 4, 5), although when tested together, the coefficient of NREM delta activity became closer to zero (Figure 2D and 3D). This suggests that NREM delta activity is likely to be beneficial in reducing wakeful neural malfunctioning in PD, which is consistent with prior research on NREM sleep and neurodegeneration (Schreiner et al. 2019; J. Chen et al. 2024; Anjum et al. 2024). However, even in the univariate models, it is not as strong a predictor as REM delta activity, at least in the motor-related neural network signatures we tested. The observed REM delta activity was unlikely to be eye-movement artifacts in our intracranial cortical data, especially recorded at a brain location distant from the orbits and using a nearest-neighbor bipolar recording configuration(Kovach et al. 2011). Beyond the existing literature, our results underscore the importance of REM sleep regarding the neural health of PD in particular and neurodegenerative diseases in general.

Furthermore, we found that the effect of REM sleep (duration and delta activity) was not constant throughout the night. Instead, its effect started from nearly zero in the middle of the sleep course and got increasingly stronger as sleep progressed (Figure 5C and 5D). This is congruent with the fact that REM sleep became more prevalent as sleep progressed (Figure 1G) and the delta activity during REM sleep also increased in strength (Figure 5B). This suggests a timing effect of REM sleep and its delta activity: REM delta activity becomes increasingly beneficial as sleep progresses, possibly due to a cumulative effect. Alternatively, this could also simply be a by-product of the increased REM delta power in later hours: REM delta activity matters more when it is stronger. In contrast to REM delta, we show that the effect of NREM delta activity, despite its generally stronger power throughout the night, was typically insignificant and held stable over the sleep course. Our finding of this timing effect of REM sleep may have implications for PD patients who typically experience early awakenings in the morning. As REM sleep gets more prevalent and its delta activity gets stronger towards morning time, these PD patients likely experience more critical loss of REM sleep and are thus more susceptible to the disease progression, which may warrant extra research attention.

### Protective role of delta activity during sleep

Our results suggest a protective role of slow-wave delta activity during sleep in relation to daytime pathophysiological neural network activity as well as subjective sleepiness symptoms in PD. Particularly in REM sleep, delta activity predicts reduced resting beta oscillation and hyper-effective connectivity in the BG-cortical network, which are neuropathological hallmarks of PD (Brown 2003). This is consistent with previous findings that delta activity during sleep is beneficial in several ways in neurodegeneration diseases (Wunderlin et al. 2020; Lee et al. 2020; Ju et al. 2017; Schreiner et al. 2019; J. Chen et al. 2024). It is worth noting that this effect of delta was not driven by the inverse relationship between delta and beta activity during sleep in PD (Anjum et al. 2024). Beta activity during sleep was numerically predictive of increased pathological neural signatures next morning but this effect was less strong than the effect of delta activity, especially when they were tested together (Figure 2D and 3D). Moreover, delta activity during REM was also associated with less sleepiness and more alertness in patients’ subjective reports. This self-report result complements and strengthens the neurophysiological assessments, as sleepiness is a common clinical symptom among PD patients (Lima 2013). Overall, this suggests that during sleep, slow-wave delta activity likely plays a central role in attenuating the pathophysiological neural activity and plausibly its associated symptoms in PD.

Beyond calculating delta power, we further detected individual delta waves in the cortical channel during nighttime recordings. Consistent with prior research (Bernardi et al. 2019), we identified delta waves in both NREM and REM stages. We show that their amplitude and occurrence rate were greater during N3 than REM sleep. Interestingly, the peak amplitude of these delta waves was a much stronger negative predictor of next-morning pathophysiological features than its occurrence rate. This suggests that the appearance of delta waves with a large amplitude may be a direct benefiting factor for neural network health in PD.

Consistent with results of the overall delta power, we also show that the LME coefficient of the delta wave peak amplitude was larger in REM sleep compared to NREM stages, underscoring the unique role of REM delta waves. Using EEG, prior research has shown that the “sawtooth” delta waves during REM sleep have a frontal-central source, matching the position of our ECoG electrodes. Unlike typical slow waves during N3, “sawtooth” delta waves in REM tend to be associated with increased eye movements and gamma activity, suggesting increased neuronal firing (Bernardi et al. 2019; Steriade, Amzica, and Contreras 1996). It is possible that REM-specific “sawtooth” waves, together with associated activities, play an important role in regulating the BG-cortical circuit in PD.

### Clinical implications for PD

Our findings regarding the beneficial role of REM sleep in PD progression carry significant clinical and translational implications. Firstly, these results suggest that PD patients suffering from sleep disturbances, particularly those with disrupted REM sleep or early awakening issues, may represent a more vulnerable subgroup at a risk of accelerated disease progression. This warrants further research validation. Secondly, the observed beneficial effects of REM sleep delta activity open novel therapeutic avenues, particularly for neuromodulation strategies. Specifically, DBS, a well-established treatment for motor symptoms in PD, could be explored for its potential to enhance overall sleep architecture. More precisely, it could be configured to promote delta activity during REM sleep, thereby leveraging the neuroprotective mechanisms of sleep in PD. Lastly, in our exploratory analyses, we observed that gamma power (broadband 30-60 Hz and entrained gamma at half-stimulation frequency) during sleep was associated with increased pathophysiological BG-cortical network features the next morning (Supplementary Figure 2, 4, 5). This observation requires future confirmatory research investigation.

## Limitations

Several limitations of our study need to be acknowledged. First, the intracranial electrodes in our participants were limited in spatial distribution and this may have favored the detection of sawtooth delta activity during REM over NREM as it relates to network health signals. Due to clinical necessity, the intracranial electrodes were implanted in the BG-motor cortex network in these PD patients, as part of a parent clinical trial of daytime adaptive DBS (Oehrn et al. 2024). As a result, it is possible that this study might have under-estimated the potential importance of NREM slow-wave activity. Second, we were able to include only four PD patients in our research protocol. However, this lower number of subjects was counteracted by utilizing the multiple-night within-subject design, enabling robust statistical LME models to account for both between- and within-subject effects. In addition, these four patients are all male in sex. It remains unclear how well our findings will generalize to other PD patients or sexes. Third, we used a portable PSG headband for sleep recording and a machine learning algorithm for sleep staging. Although this provides self-implemented continuous recording at home and fast automated scoring, it may have lower accuracy compared to a full set of PSG scored by certified clinicians (González et al. 2024; Ravindran et al. 2025). However, we do not believe that this will have confounded our REM sleep results as there were clear separations in the power spectrum between REM vs. awake and REM vs. N2 in our data (Supplementary Figure 1A), where misclassifications are most likely to occur. Finally, our participants included mixed locations of subcortical electrodes, which may have blocked a detection of a sleep effect on the early EP amplitude, an index of antidromic activation of the hyperdirect pathway. Nevertheless, we found sleep effects on EP attributes in later windows despite the mixed subcortical sites, suggesting a robustness of the findings on the BG-cortical orthodromic pathway.

### Conclusion

We show that nighttime REM sleep predicts reduced abnormal daytime activities in a key neural network implicated in PD, by following a group of PD patients for multiple nights using wireless multi-site intracranial recording. Specifically, we assessed multiple established objective pathophysiological BG-cortical network signatures associated with PD symptoms and progression, plus subjective self-reports. We found that longer and faster onset of REM sleep was associated with a reduction in these pathophysiological signatures. We further show that delta activity during REM sleep plays a central role in this beneficial effect in attenuating pathophysiological activities and improving subjective alertness after sleep. Our findings highlight a potentially protective role of REM sleep in the progression of Parkinson’s disease. Future research is needed to further examine the role of REM sleep in other neurodegenerative diseases and other aspects of disease progression.

## Methods

### Participants

The study was registered on clinicaltrials.gov (NCT0358289; IDE G180097), and the study protocol was approved by the Institutional Review Board of the University of California, San Francisco. The four participants presented in this study were diagnosed with idiopathic Parkinson’s disease (Table 1), and were enrolled as part of a parent study investigating daytime adaptive deep brain stimulation protocols (Oehrn et al. 2024).

**Table 1.**
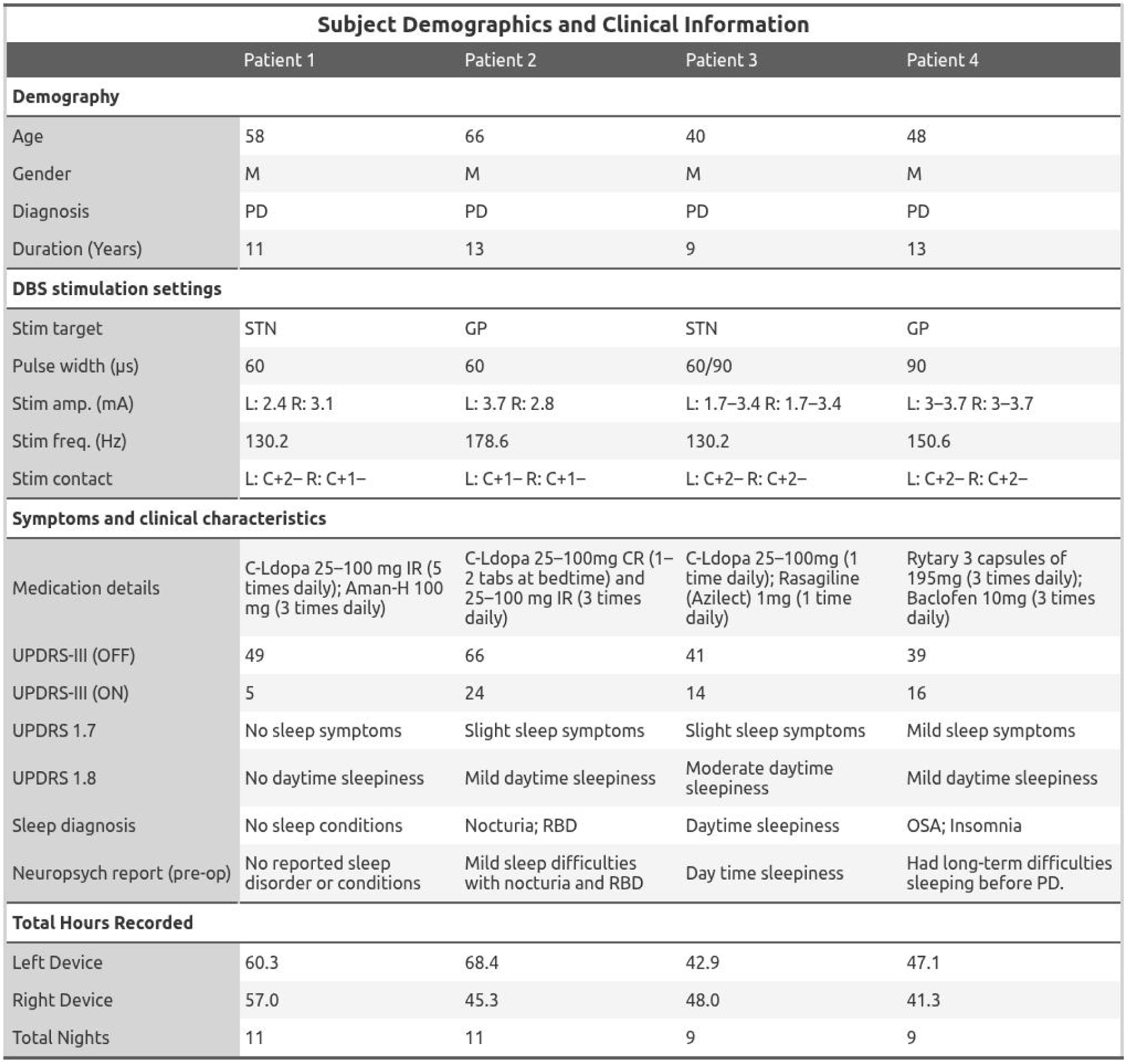
Subject Information. . *C-Ldopa: Carboxy-Levodopa. Aman-H: Amantadine-Hydrochloride. RBD: REM-sleep Behavior Disorder. OSA: Obstructive Sleep Apnea. UPDRS: Unified Parkinson’s Disease Rating Scale (obtained prior to DBS implantation)*

| Subject Demographics and Clinical Information |  |  |  |  |
| --- | --- | --- | --- | --- |
|  | Patient 1 | Patient 2 | Patient 3 | Patient 4 |
| **Demography** |  |  |  |  |
| Age | 58 | 66 | 40 | 48 |
| Gender | M | M | M | M |
| Diagnosis | PD | PD | PD | PD |
| Duration (Years) | 11 | 13 | 9 | 13 |
| **DBS stimulation settings** |  |  |  |  |
| Stim target | STN | GP | STN | GP |
| Pulse width (μs) | 60 | 60 | 60/90 | 90 |
| Stim amp. (mA) | L: 2.4 R: 3.1 | L: 3.7 R: 2.8 | L: 1.7–3.4 R: 1.7–3.4 | L: 3–3.7 R: 3–3.7 |
| Stim freq. (Hz) | 130.2 | 178.6 | 130.2 | 150.6 |
| Stim contact | L: C+2– R: C+1– | L: C+1– R: C+1– | L: C+2– R: C+2– | L: C+2– R: C+2– |
| **Symptoms and clinical characteristics** |  |  |  |  |
| Medication details | C-Ldopa 25–100 mg IR (5 times daily); Aman-H 100 mg (3 times daily) | C-Ldopa 25–100mg CR (1–2 tabs at bedtime) and 25–100 mg IR (3 times daily) | C-Ldopa 25–100mg (1 time daily); Rasagiline (Azilect) 1mg (1 time daily) | Rytary 3 capsules of 195mg (3 times daily); Baclofen 10mg (3 times daily) |
| UPDRS-III (OFF) | 49 | 66 | 41 | 39 |
| UPDRS-III (ON) | 5 | 24 | 14 | 16 |
| UPDRS 1.7 | No sleep symptoms | Slight sleep symptoms | Slight sleep symptoms | Mild sleep symptoms |
| UPDRS 1.8 | No daytime sleepiness | Mild daytime sleepiness | Moderate daytime sleepiness | Mild daytime sleepiness |
| Sleep diagnosis | No sleep conditions | Nocturia; RBD | Daytime sleepiness | OSA; Insomnia |
| Neuropsych report (pre-op) | No reported sleep disorder or conditions | Mild sleep difficulties with nocturia and RBD | Day time sleepiness | Had long-term difficulties sleeping before PD. |
| **Total Hours Recorded** |  |  |  |  |
| Left Device | 60.3 | 68.4 | 42.9 | 47.1 |
| Right Device | 57.0 | 45.3 | 48.0 | 41.3 |
| Total Nights | 11 | 11 | 9 | 9 |

Participants were implanted with bilateral quadripolar subcortical electrodes capable of both sensing and stimulation. Patient 1 and 3 received these electrodes in the Subthalamic Nucleus (STN) using Medtronic model 3389 leads, and Patient 2 and 4 were implanted in the Globus Pallidus (GP) with Medtronic model 3387 leads. The selection of the implant site was determined by the clinical team of each participant according to standard clinical criteria. Subdural quadripolar electrocorticogram (ECoG) arrays (Medtronic Resume II, model 0913025) were implanted bilaterally over the central sulcus. All electrodes were connected to Summit RC+S (Medtronic model B35300R) implantable neural stimulators (INS), which have been extensively described in previous studies (Smyth et al. 2023; Anjum et al. 2024; Gilron et al. 2021; Oehrn et al. 2024). The Summit RC+S system enables the recording of field potentials from both cortical and basal ganglia locations during chronic stimulation. It also allows for the streaming of data recordings to a nearby tablet for offline analysis while participants are at home.

### Experimental procedures

Participants maintained their typical sleep schedule at home for approximately 10 nights, most of which were consecutive. Overnight sleep at home was recorded concurrently using portable PSG via the DREEM2 headset, alongside intracranial BG-cortical data streaming via the RC+S system. After awakening and before medication use, participants had a morning assessment session that included self-report questionnaires, 1-minute resting-state recording, and 4-minute evoked potential recording (2 minutes each hemisphere). Portable PSG was removed upon awakening, and only intracranial data were recorded in the morning sessions. All data recordings were conducted remotely in the participants’ homes. All participants continued their regular dopaminergic medication throughout the study. During sleep and resting-state recording, participants received their clinical continuous DBS therapy and regular medications. We initially collected a total of 42 night+morning pairs of data. One morning session data of Patient 3 and Patient 4 failed to be recorded and were thus excluded. Valid data of a total of 40 night+morning pairs were used for analysis.

### Overnight data acquisition

The DREEM2 headband collects electroencephalography (EEG) data and additional signals including pulse oximetry, electromyography, and accelerometry. The DREEM2 headband provides sleep stage classification hypnograms with an automated sleep staging algorithm, trained to emulate AASM scoring methods. The accuracy of these automatic sleep stage labels is comparable to that provided by certified experts (Arnal et al. 2020). Validations of the DREEM2 sleep staging in older adults (some with PD and AD) also showed good accuracy, ranging from 80-93% for REM, 70-89% for N1/N2, and 93-95% for N3 (González et al. 2024; Ravindran et al. 2025). This has also been previously validated as matching expected intracranial neurophysiology in PD by our group in a separate study focused on NREM sleep (Anjum et al. 2024).

Three intracranial field potential streams were recorded at a sampling rate of 500 Hz. The basal ganglia potentials were measured using a bipolar ‘sandwich’ electrode configuration that captures the electric potential difference between two electrodes adjacent to the stimulation electrode. Cortical field potentials were recorded in two streams: one using a bipolar configuration on the precentral gyrus and another bipolar configuration spanning the entire electrode array across the central sulcus. Participants recorded about ten nights of both external polysomnogram and intracranial field potential data during ongoing therapeutic stimulation.

Data alignment involved correlating sleep stages from the polysomnogram with intracranial field potentials. To address discrepancies in sampling rates and time offsets between the DREEM and RC+S data streams, cross-correlation analysis was employed on the accelerometry data from each device, adjusting the temporal alignment to maximize correlation, as detailed in our previous report (Anjum et al. 2024). The resulting dataset includes a sleep stage label (N1, N2, N3, REM, Wake) for each 30-second epoch of the intracranial field potential streams for each hemisphere.

### Morning data acquisition

During resting-state morning recordings, intracranial data were streamed with the same settings as the overnight recordings. Participants indicated they awoke via a checkbox on their streaming application. Participants completed self-report questionnaires on their previous sleep and current subjective state. Specifically, participants reported their sleepiness level on a 1-9 Likert scale using the Karolinska Sleepiness Scale (Kaida et al. 2006), and their ‘restedness’ level from sleep on a 1-5 scale using the Consensus Sleep Diary (Carney et al. 2012). Following that, participants recorded neural activity during wakeful quiescence. Participants subsequently self-triggered their evoked potential procedure, which automatically and programmatically cycled through the evoked potential stimulation paradigm. During the evoked potential paradigm, stimulation pulses were delivered in each hemisphere at the basal ganglia stimulation electrode at a rate of 4 Hz, with an amplitude of 4 mA, and with the same pulse width as their clinical stimulation (60 or 90 µs). Field potentials were recorded at a sampling rate of 1000 Hz from one cortical electrode and one subcortical electrode ipsilaterally. This procedure was implemented for 2 minutes in each hemisphere at a time, with the contralateral DBS off (e.g. left device for 2 minutes with right device off, then vice versa).

### Overnight intracranial data analysis

#### Power spectrum analysis

Power spectra were calculated in 2-second windows for each participant and each sleep stage via Welch’s technique using the scipy python package, with a Hann window, 1000-point segments (2s sample), and 0 overlap between segments. Delta power was the sum of power from 0.5-4 Hz and beta power was the sum of power from 13.5-30 Hz. The power of the two cortical streams was averaged. Both delta and beta powers were log-transformed, and then normalized with z-scoring across all nights within each hemisphere of each patient before statistical tests.

#### Detection of delta waves

From the intracranial cortical data during overnight sleep recordings, we detected delta waves using a previously established method (Bernardi et al. 2019; Riedner et al. 2007). Specifically, the signal of each cortical channel was first down-sampled to 125 Hz and bandpass filtered between 1 and 10 Hz (the upper limit of the filter was selected to minimize wave shape and amplitude distortions). An algorithm based on detecting consecutive signal zero-crossings (Bernardi et al. 2019; Riedner et al. 2007) was used to detect delta waves during sleep. Negative half-waves with a duration of consecutive zero-crossings between 125 and 500 ms (1– 4 Hz) were extracted, and the following properties were quantified: peak amplitude (mV, in absolute value) and duration (ms). The average peak amplitude and occurrence rate were quantified in each sleep stage. Raw signal before down-samping and filtering corresponding to the detected negative half-waves was used for delta wave visualization (Figure 5A).

#### Detection of beta bursts

The intracranial cortical data were first down-sampled to 250 Hz and bandpass filtered in the beta frequency range (13-30 Hz). The amplitude envelope was calculated on the preprocessed data using a Hilbert transform (‘hilbert’ from scipy). For each channel across all the nights of a participant, the 75th percentile of the beta amplitude envelope was set as the threshold of a beta burst, following prior research (Tinkhauser et al. 2017). Beta bursts were detected as the amplitude envelope exceeding the threshold, and their average peak amplitude and occurrence rate were calculated in each sleep stage.

### Morning resting-state data analysis

Intracranial data of all channels in both hemispheres during the resting-state recording (1 min) were extracted for each morning session. The power spectral density was calculated for the extracted data using Welch’s method (“psd_array_welch” in MNE time-frequency) with a Hamming window, 500-point segments (1s sample), and 0 overlap. Beta band power was calculated as the sum of power from 13-30 Hz and further log-transformed for later analyses.

The BG-cortical functional connectivity was quantified as the coherence between the basal ganglia channel and the cortical channels in the extracted resting-state data, with a “cwt_morlet” time-frequency decomposition method and a hanning smoothing kernel (“spectral_connectivity_time” in MNE_connectivity). Coherence at the beta frequency (13-30 Hz) was extracted and averaged across frequency bins. As LME models on BG-cortical functional connectivity outcomes had singular fits with ‘hemisphere’ as a random effect, ‘hemisphere’ was dropped as a random effect here.

### Evoked potential data analysis

Intracranial data of subcortical and cortical channels during the evoked potential recording were extracted for each evoked response into 250-ms epochs (“mne.EpochsArray”) with 50ms pre-stimulation baseline. For each recording session, the data from the beginning 10s and the ending 2s were excluded from analyses because of signal disruption due to changes of stimulation frequency and amplitude. Each epoch’s data were baseline corrected and epochs were rejected if the amplitude in the cortical channel exceeded 200 µV or in the subcortical channel exceeded 1000 µV. Temporally misaligned epochs due to packet loss in the RC+S system were visually inspected and rejected. After preprocessing, each hemisphere retained an average of 491 (SD=146, median=468) evoked responses in a session (see Figure 2A and 2B for example plot of epoch-level and session-level EP respectively).

For each EP epoch of the cortical channel data, we calculated the peak-to-trough amplitudes in an early window (0 to 10 ms) and a main window (10 to 100 ms), and the power spectral density of the whole evoked response excluding the early window (10-200 ms) using Welch’s method with a Hamming window (“psd_array_welch” in MNE time-frequency). Beta band power was calculated as the sum of power from 13-30 Hz. These epoch-level EP metrics were averaged for each channel during each session, and then log-transformed for later analyses.

### Statistical tests

We used Linear Mixed-Effects (LME) Models to test the effects of sleep parameters on next-morning pathophysiological neural signatures in R, with the “lmer” function from “lme4” and “lmerTest” packages. The coefficients were estimated using the Maximum likelihood method. In the main analyses, we include both REM and NREM parameters in the same model as the fixed effects, and participant and hemisphere as nested random effects on intercept. Specifically, we tested the effects of REM and NREM sleep duration or latency together in the “duration” models. As sleep stage duration and latency were anti-related and would introduce colinearity issue in the same LME model, we tested the duration predictors in one model (outcome ∼ NREM_duration + REM_duration + (1|subject/hemisphere)) and the latency predictors in another (outcome ∼ NREM_latency + REM_latency + (1|subject/hemisphere)). We tested the effects of beta and delta power in REM and NREM together in the “power” models (outcome ∼NREM_beta + REM_beta + NREM_delta + REM_delta + (1|subject/hemisphere)). In the secondary analyses testing the effects of delta power and sleep stage percentage over time, we tested the effects of individual predictors in separate LME models and obtained their model coefficients with 95% confidence intervals. In the subsequent analyses of testing the effects of delta power vs. delta wave metrics and beta power vs. beta burst metrics, we tested the effects of individual predictors in separate LME models and obtained their standardized statistics (t-stat) because the predictors are on different scales. In the prediction of self-report measures, as each session had only one report per participant (as opposed to per hemisphere in the neurophysiological outcomes), the random effect thus included only a random intercept of ‘participant’.

## Supplementary Materials

**Supplementary Figure 1.**
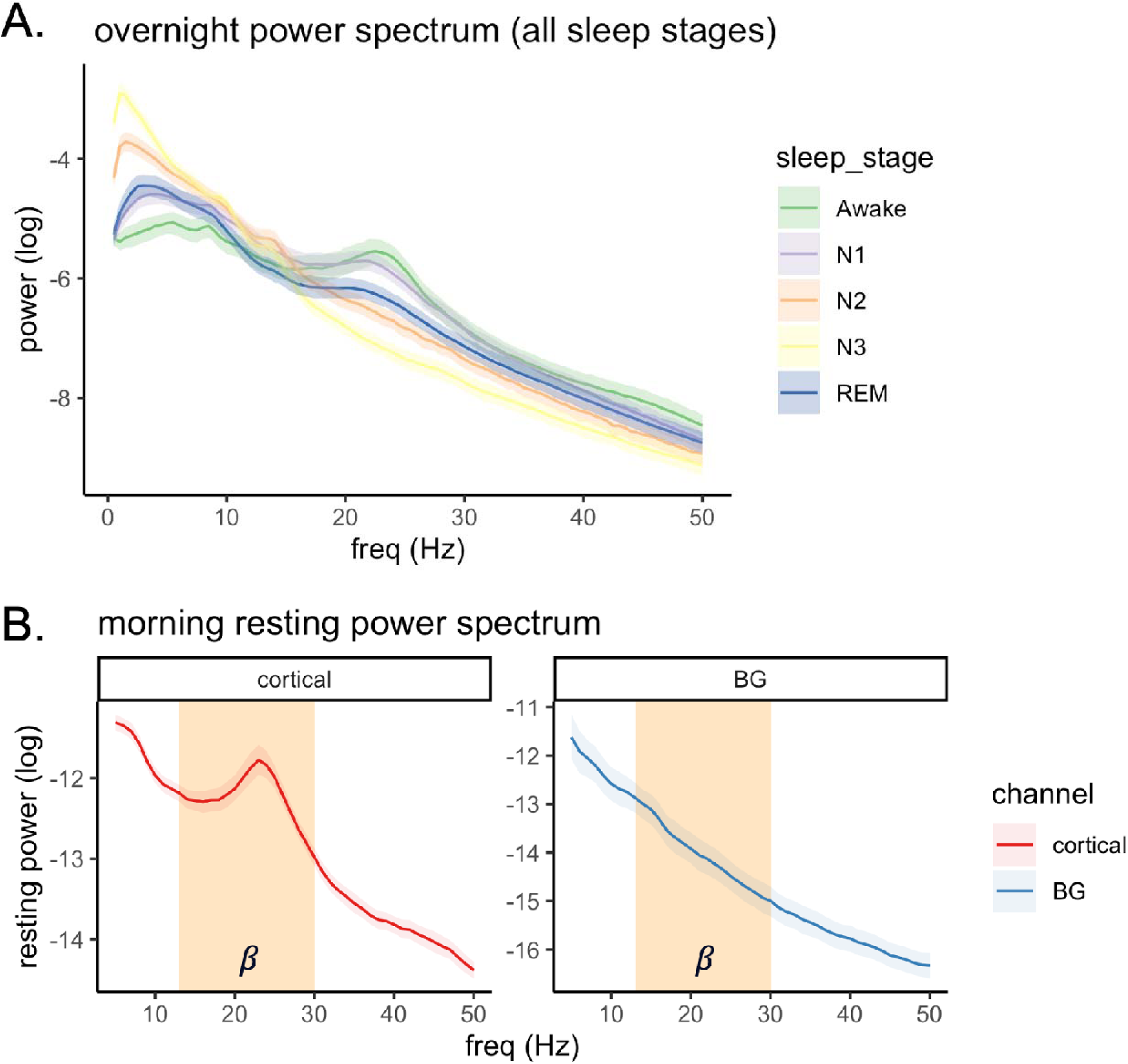
Overnight and morning power spectrum in all subjects. (A) Power spectrum of overnight intracranial cortical recordings in different sleep stages. (B) Power spectrum of morning resting recording in the cortical vs. basal ganglia (BG) channels. There is a power peak in the beta frequency (13-30 Hz, highlighted) in the cortical channels and not in the BG channel. Shading areas indicate 95% confidence intervals across days.

**Supplementary Figure 2.**
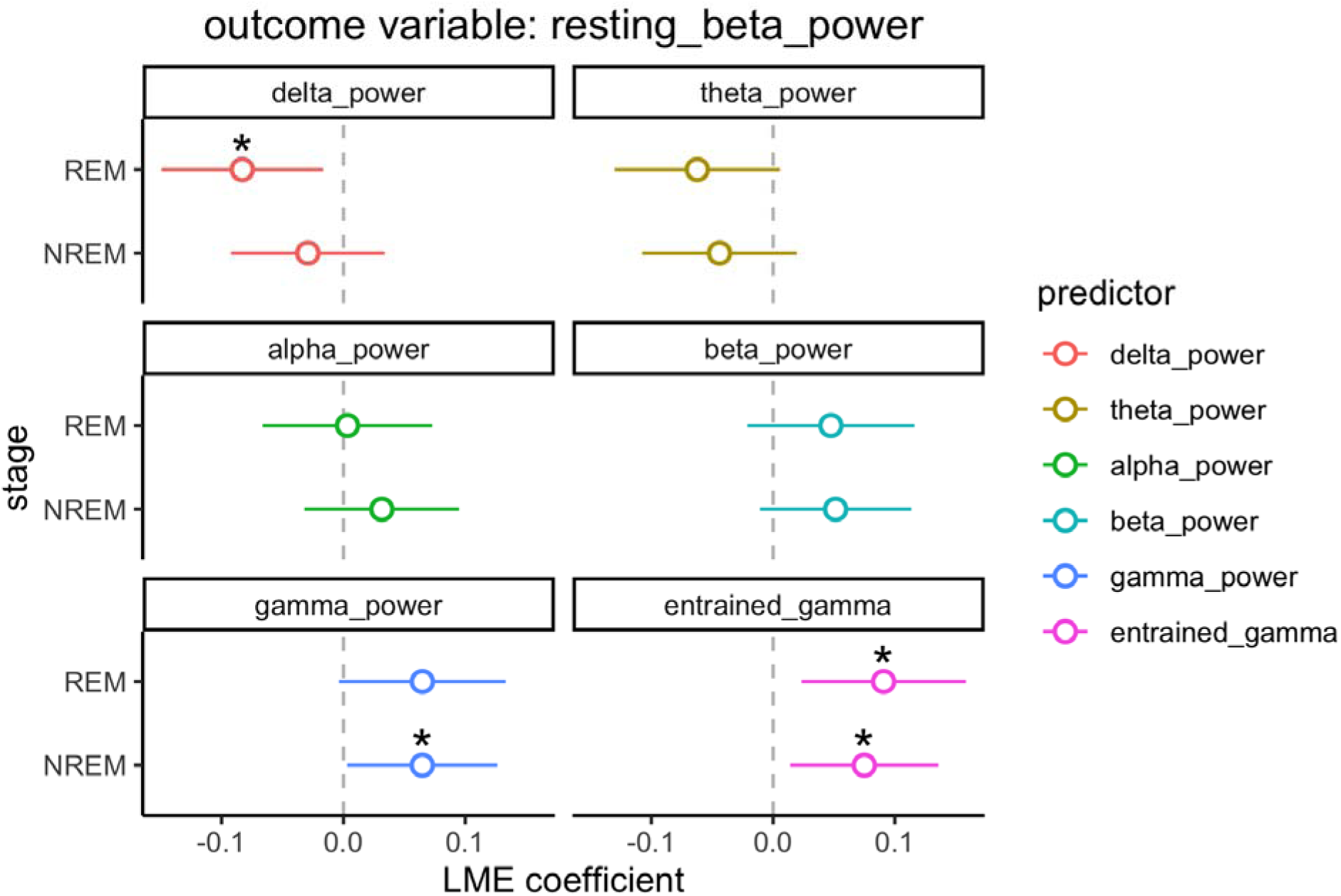
The results of exploratory linear mixed-effect (LME) models of multiple spectral power signals in prediction of resting-state beta power. The power predictors include delta power (1-4 Hz), theta power (4-8 Hz), alpha power (8-13 Hz), beta power (13-30 Hz), gamma power (30-60 Hz), and entrained gamma power (±2 Hz around patients’ half-stimulation frequency). Each of these power predictors was tested in individual LME models separately, and also separately for REM and NREM power. Error bars indicate 95% confidence intervals of LME coefficient. Asterisks indicate statistical significance in LME models: * = *p*<.05, ** = *p*< .01, *** = *p*< .001.

**Supplementary Figure 3.**
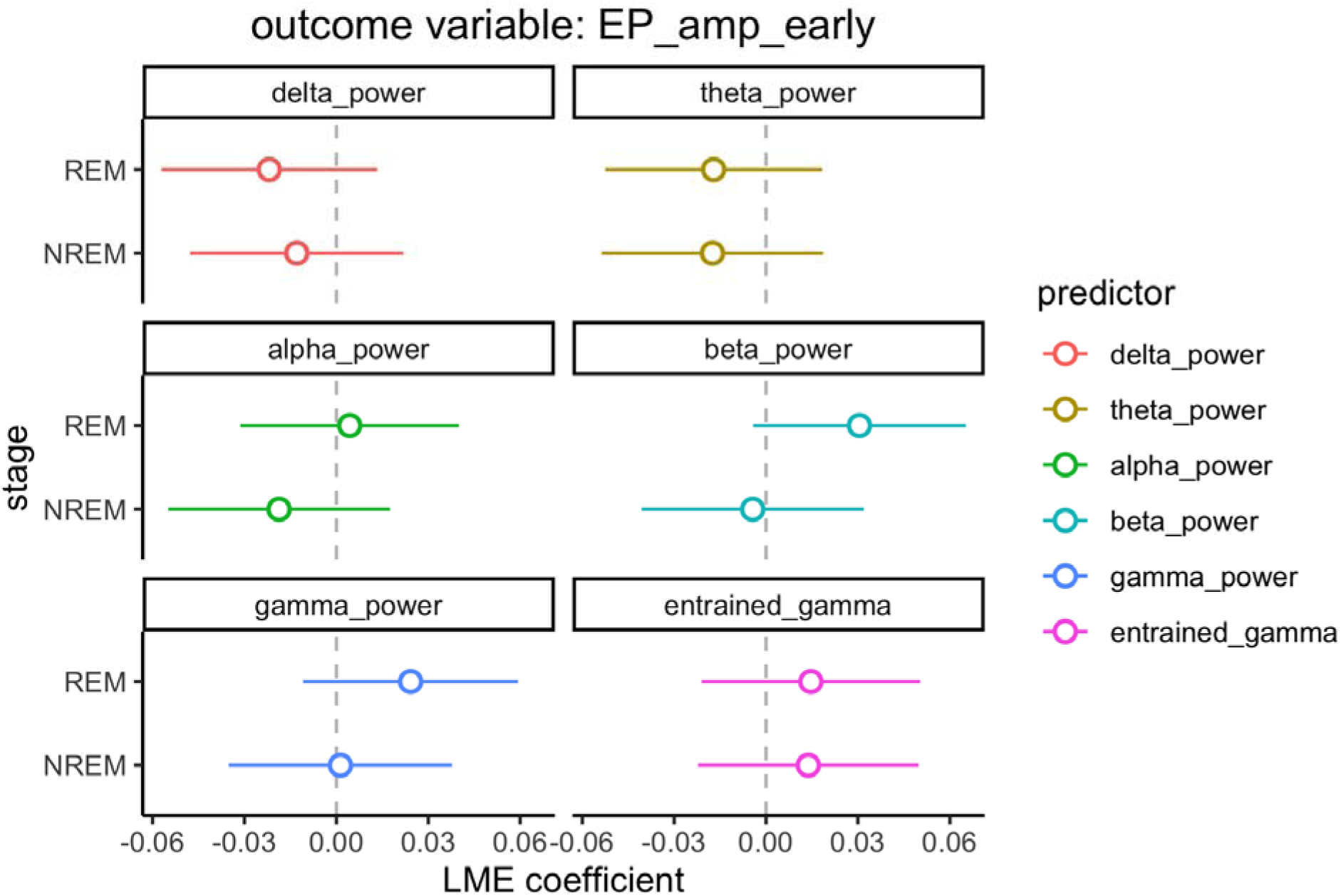
The results of exploratory linear mixed-effect (LME) models of multiple spectral power signals in prediction of amplitude of evoked potentials (EP) in the early window. The power predictors include delta power (1-4 Hz), theta power (4-8 Hz), alpha power (8-13 Hz), beta power (13-30 Hz), gamma power (30-60 Hz), and entrained gamma power (±2 Hz around patients’ half-stimulation frequency). Each of these power predictors was tested in individual LME models separately, and also separately for REM and NREM power. Error bars indicate 95% confidence intervals of LME coefficient. Asterisks indicate statistical significance in LME models: * = *p*<.05, ** = *p*< .01, *** = *p*< .001.

**Supplementary Figure 4.**
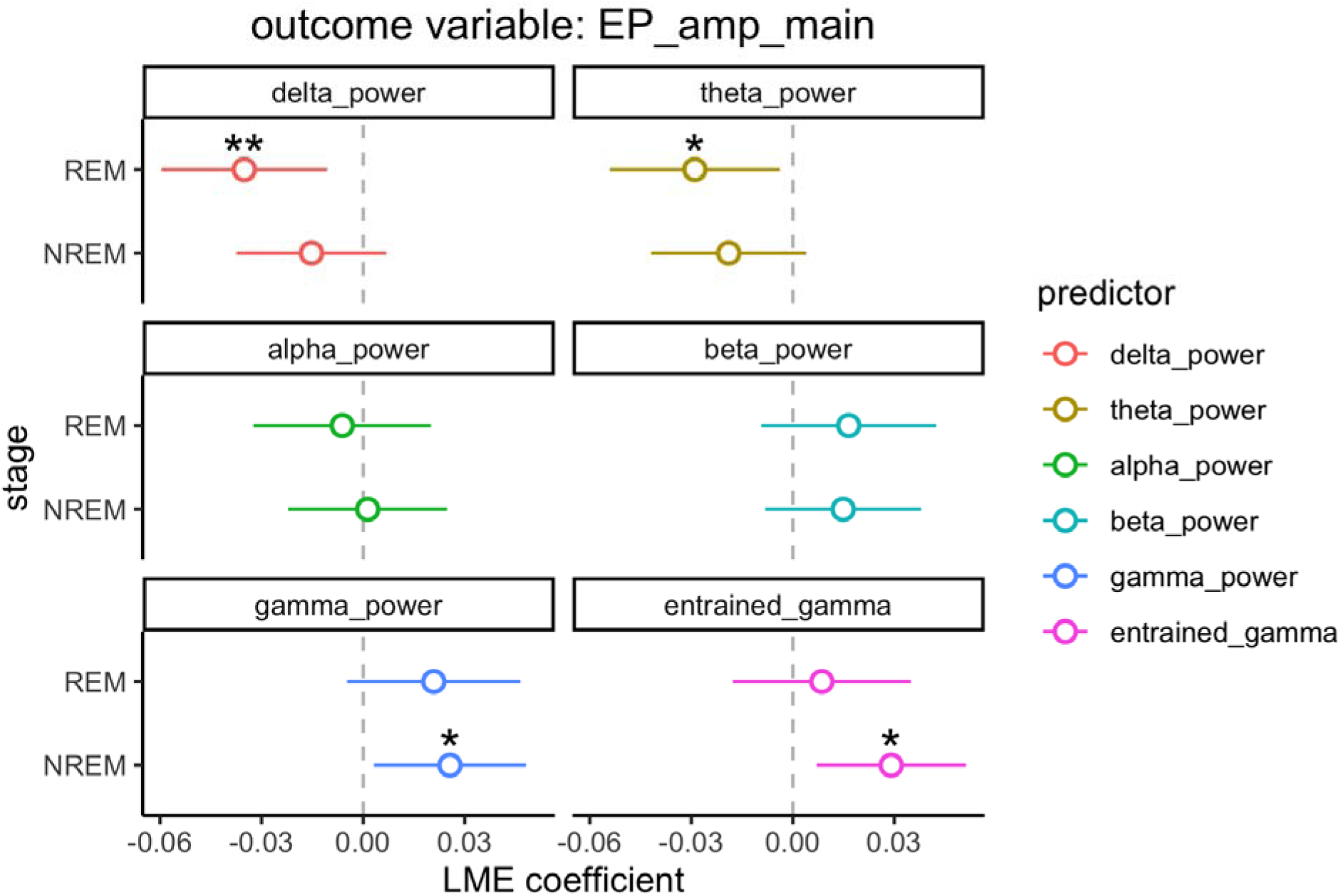
The results of exploratory linear mixed-effect (LME) models of multiple spectral power signals in prediction of amplitude of evoked potentials (EP) in the main window. The power predictors include delta power (1-4 Hz), theta power (4-8 Hz), alpha power (8-13 Hz), beta power (13-30 Hz), gamma power (30-60 Hz), and entrained gamma power (±2 Hz around patients’ half-stimulation frequency). Each of these power predictors was tested in individual LME models separately, and also separately for REM and NREM power. Error bars indicate 95% confidence intervals of LME coefficient. Asterisks indicate statistical significance in LME models: * = *p*<.05, ** = *p*< .01, *** = *p*< .001.

**Supplementary Figure 5.**
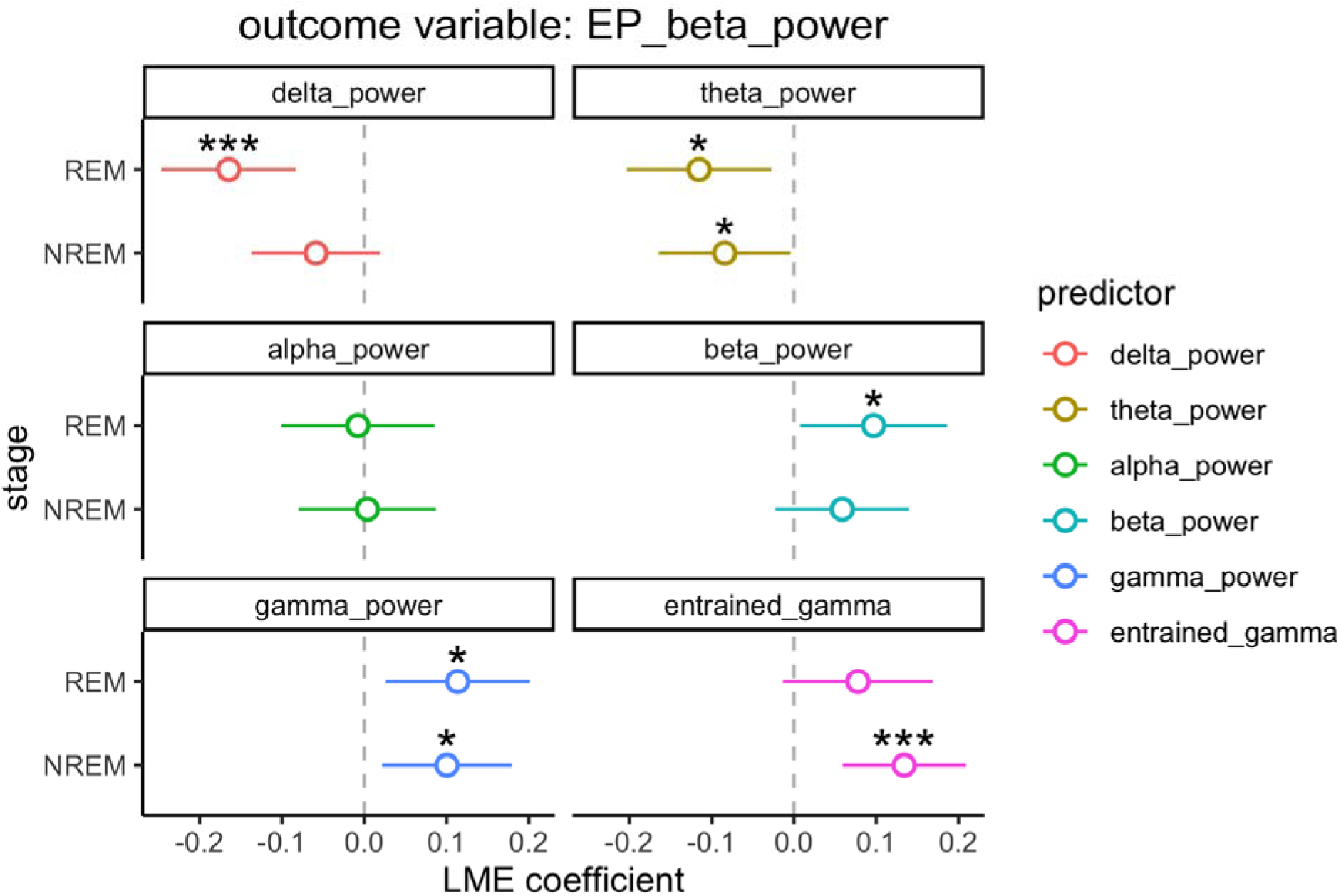
The results of exploratory linear mixed-effect (LME) models of multiple spectral power signals in prediction of the beta power of evoked potentials (EP). The power predictors include delta power (1-4 Hz), theta power (4-8 Hz), alpha power (8-13 Hz), beta power (13-30 Hz), gamma power (30-60 Hz), and entrained gamma power (±2 Hz around patients’ half-stimulation frequency). Each of these power predictors was tested in individual LME models separately, and also separately for REM and NREM power. Error bars indicate 95% confidence intervals of LME coefficient. Asterisks indicate statistical significance in LME models: * = *p*<.05, ** = *p*< .01, *** = *p*< .001.

